# Elucidating the link between binding statistics and Shannon information in biological cooperative networks

**DOI:** 10.1101/2024.03.28.587158

**Authors:** Kinshuk Banerjee, Biswajit Das

**Author notes:** These authors contributed equally to this work.

## Abstract

Cooperative response is ubiquitous and vital for regulatory control and ultra-sensitivity in various cellular biophysical processes. Ligands, acting as signaling molecules, carry information which is transmitted through the elements of the biochemical network during binding processes. In this work, we address a fundamental issue regarding the link between the information content of the various states of the binding network and the observable binding statistics. Two seminal models of cooperativity, *viz*., the Koshland-Nemethy-Filmer (KNF) network and the Monod-Wyman-Changeux (MWC) network are considered for this purpsoe which are solved using the chemical master equation approach. Our results establish that the variation of Shannon information associated with the network states has a generic form related to the average binding number. Further, the logarithmic sensitivity of the slope of Shannon information is shown to be related to the Hill slope in terms of the variance of the binding number distributions.

^1^

## 1 Introduction

In living organisms, cellular communication is essential for the cellular decision-making process [1–9]. Such communication is established by transmitting the information either from one cell to another by following the inter-cellular signaling-network[9–12] or by the communication among the cellular components inside a cell through the intra-cellular signaling-pathway [13–18]. Cellular information is generally transmitted by the signaling molecules, called ligands, which bind to the receptor proteins on the cell surface [16–18]. Different types of inter-cellular communication networks, *e*.*g*., endocrine, paracrine, synaptic, and autocrine are observed in mammalians which regulate the growth, division, as well as organization of the cells[17, 18]. For example, the power-house of the cell, mitochondria in different tissues talk to each other through the inter-signaling network to repair the injured cells. When their signal fails, the biological clock starts winding down; and consequently, cell-aging is started[11]. The transmitted information through this network can be measured nowadays from the single-cell RNA-sequencing (scRNA-seq) data, known as CellChat[10]. Similarly, binding of the signaling molecules to the receptors affects the intra-cellular communication network which changes the intra-cellular physiology[17, 18], *e*.*g*., the mitogen activated protein (MAP)-kinase signaling network where MAP-kinase phosphorylates the transcription factors to transcript some specific genes[19, 20] and non-genomic calcium signaling network which changes the intra-cellular calcium concentration[21, 22].

Inter- and intra-cellular signaling networks exhibit significant levels of noise in the response of identical cells in the same cellular environment as the transmission of information varies from cell to cell due to the variation in the binding affinity of the signaling molecules to the receptors at the molecular scale[1–6]. This variation in the binding affinity of ligands leads to a cooperative and collective response of the cell or cell cluster [3–5]. For example, delayed sigmoidal response of the cells is a signature of the cooperative enhancement of the binding tendency of the signaling molecules to the cell receptors[23–25]. The cooperative response is not only observed in the inter- or intra-cellular networks but also in oligomeric or multimeric proteins and enzymes having more than one ligand-binding subunits [26–33]. The classical example is haemoglobin, an oligomeric protein consisting of four ligand-binding subunits, exhibiting sigmoidal response during oxygen binding[30, 33]. Communication among these subunits alter the ligand-binding affinity in successive binding steps which leads to cooperative response [28, 29, 31, 32]. Actually, the long-range communication among the amino acids in a protein, called allosteric communication, leads to such cooperativity[34, 35]. Sometimes, communication network can be formed among the protein or ligand molecules for which cooperative response can be observed *e*.*g*., zero-order ultra sensitivity of the phosphorylation-dephosphorylation cycle (PDPC)[36, 37]. An enzyme with a single subunit can also exhibit cooperativity if the communication network is formed with the slow fluctuating conformations of the protein[38]. Also, constraint in the binding process, *e*.*g*., in a channel geometry[39], can give rise to cooperative behavior.

Generally, cooperative response of a ligand-binding network is characterized in terms of the Hill coefficient which is determined from suitably-defined slopes of experimental binding isotherms called Hill plots[31, 40–44]. Hill-coefficient, the value of Hill slope at the half-saturation point, greater than unity indicates positive cooperative response where the successive communication or interaction increases the binding affinity from one element to another in a bio-physical system[31, 40, 41]. Similarly, for the reverse case, Hill coefficient becomes less than unity indicating negative cooperative response[31, 40, 41]. If the interaction among the elements remains same throughout the binding process, no cooperativity is observed in the response of the system [31, 40]. For example, in the metastasis of the cancer cells, sigmoidal response is exhibited by the cells during the colonization as the cooperative communication is established between the invading cancer cells with the tumour micro-environment[45]. However, membrane interaction of bound ligands contributes to the negative cooperative response of the EGF receptors[46, 47]. Hill coefficient is determined experimentally in terms of the average binding number of ligands[40, 42, 48]. Theoretically, the experimental binding curves are modelled with two seminal cooperative networks *viz*., the Koshland-Nemethy-Filmer (KNF) network[49] and the Monod-Wyman-Changeux (MWC) network[50].

In this study, our goal is to quantify the binding in terms of information associated with the various ligand-bound states or elements of these networks. To this end, we treat the networks using the tools of stochastic kinetics with chemical master equations (CMEs)[51] and their solutions in equilibrium. The resulting probability distributions of the binding number is then employed to characterize the information content of each bound-state in terms of the Shannon information[52]. Then, we ask the fundamental question: *Is the binding statistics, specifically the mean and variance, are related to this information, and if yes, then how?* Our work reveals that there is a clear link between the logarithmic sensitivity of Shannon information and binding statistics and the information content can be realized in terms of the experimentally measurable average binding and Hill slope.

The paper is organized as follows. In Sec.2, we give the CMEs for the different cooperative binding schemes mentioned above, along with their equilibrium solutions. The variation of Shannon information with ligand concentration and its relation to binding statistics are discussed in Sec.3. Sec.4 is devoted to numerical analyses to corroborate the analytical findings. The paper is concluded in Sec.5.

## 2 The CMEs for various binding schemes

Here we consider that a biophysical system exhibits its cooperative response either through the linear like Koshland-Nemethy-Filmer (KNF) network [49] or the ladder like Monod-Wyman-Changeux (MWC) network [50]. The response of the system is exhibited due to binding of the *n*_*T*_ number of ligand molecules to the same number of elements in the network where elements would be the receptors, or sub-units of a multimeric proteins, or the number of proteins, or the conformations of a single protein/enzyme. At cellular level, the number of elements of a network are very small compared to the number of ligand molecules. Thus, here we consider that the concentration of the ligand remain essentially constant. Suppose, there are two elements in a network, *e*_0_ and *e*_1_, where *e*_0_ is the element when no ligand is attached whereas *e*_1_ indicates the element with ligand. Dynamics of the network can be expressed in terms of the network scheme,

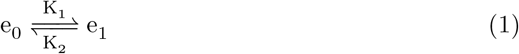

where 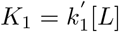 is the pseudo-first-order rate constant of binding for constant ligand concentration [*L*]. The scheme can be easily mapped with the binding scheme for a substrate *S* with a single molecule of a monomeric enzyme. Now, if there are *n*_*T*_ number of elements in the network, the scheme in (1) can be straightforwardly extended to the following with state-dependent binding and dissociation rate constants of the ligands, 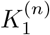 and 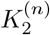, respectively:

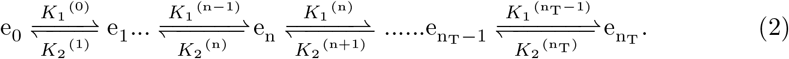

In the scheme in (2), *e*_*n*_ represents the element in a reaction network where ‘*n*’ number of ligand molecules are attached. Shifting of one to another element occurs due to binding of a ligand. Basically, ‘*n*’ is a random variable quantity and the dynamics of the pathway can be studied using a chemical master equation (CME)[51]. Using this technique, we explore the following two seminal models of cooperativity *along with a variant*:

### 2.1 The KNF network and its variant

In the Koshland-Nemethy-Filmer (KNF) network[49], the ligand molecules bind or dissociate from the element of a network sequentially with state-dependent binding constants. We also study a variant of this network where binding occurs in a channel geometry[39]. Here, the ligand molecules attach to an element only when the previous elements are occupied.and detachment occurs from the last element. This makes the network different from the KNF network where ligand molecules can bind at and detach from any element independently. The element *e*_*n*_ in this network is thus adjacent in space as well as in time and the resulting cooperativity is denoted spatial (SP)[31].

The KNF network is shown in Fig.1(a) along with that of the SP network in Fig.1(b). The corresponding CMEs can be generally formulated as

**Fig. 1.**
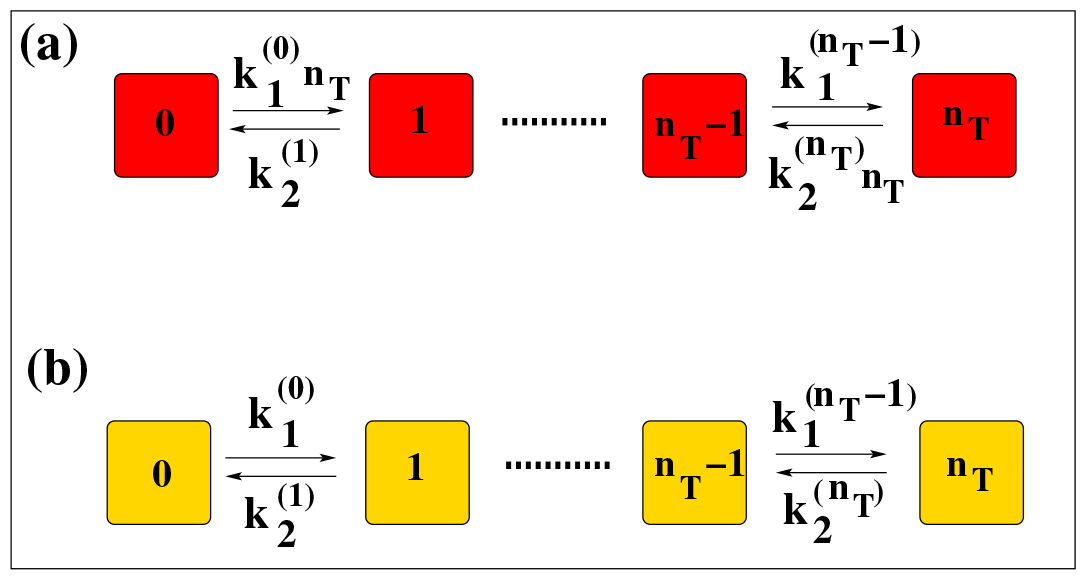
Schematic diagram of the (a) KNF network and the (b) SP network. The squares denote states of the element bound with a certain number of ligand molecules as indicated inside each square. The transition probabilities in forward and backward directions are also given (see text).

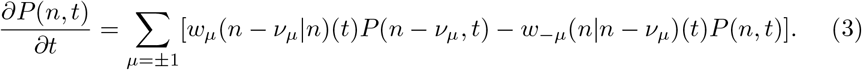

Here *μ* is the reaction index with *μ* = 1 for the forward reaction and *μ* = −1 for the reverse reaction (see the scheme in (2)). The stoichiometric coefficients *ν*_*μ*_ have values *ν*_1_ = 1 and *ν*_−1_ = −1. The probability of having ‘*n*’ number of elements being occupied by ligand molecules at time *t* is denoted by *P*(*n, t*). Here *n* runs from 0 to *n*_*T*_, the total number of elements in the network. The transition probabilities between the elements, *w*_*μ*_, are defined for the KNF network by

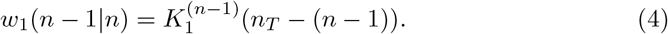

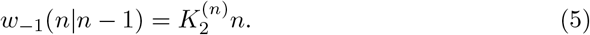

For the SP network, they are written as

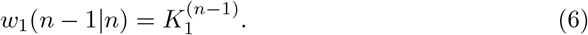

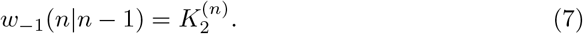

For clarity, we label *P*(*n, t*) by *P*_KNF_(*n, t*) for the KNF network and by *P*_SP_(*n, t*) for the SP network. The equilibrium solutions (denoted with superscript ‘e’) of the CMEs for the above networks are obtained by setting the right-hand-side of the set of equations, Eq.3, equal to zero. They are given below:

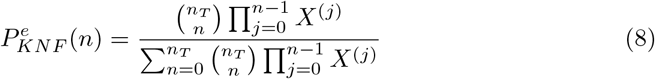

and

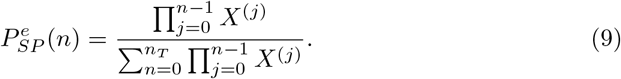

We define the state-dependent binding constants as

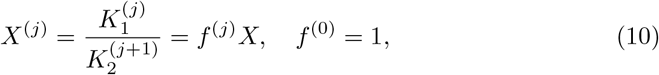

where

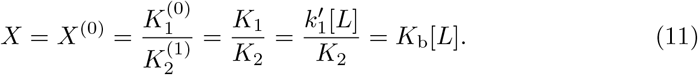

For *X*^(*j*)^ = *X* ∀*j*, the KNF network reduces to a non-cooperative (NC) network with a binomial distribution

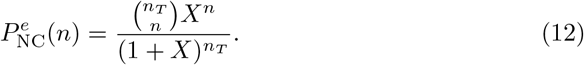

The mean binding number is defined as ⟨*n*⟩ = Σ_*n*_ *nP*^*e*^(*n*). However, it can be analytically determined in a closed form only for the NC scheme given by

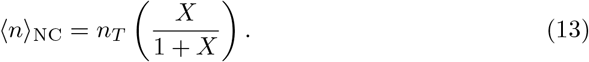

We may mention here that, for *n*_*T*_ > 1, the SP network shows cooperative behavior even for state-independent binding constants[31].

### 2.2 The MWC network

Following the Monod-Wyman-Changeux (MWC) network,[50] here we consider that an element in this reaction network resides either in ‘active’ state, represented as *R* or in ‘inactive’ state, designated here as *T*. R and T have different ligand-binding affinities. The ‘R’-state has a greater ligand-binding affinity compare to T. Initially, ligand molecules bind to the ‘T’-state, however, the population of R-states gradually increases with the progress of the reaction. The MWC network is shown schematically in Fig.2. The dynamics of the network can be described in terms of the number of elements bound by the ligands with in ‘R’ or ‘T’-states. Here, *K*_1,C_ and *K*_2,C_ (C = R, T) are the forward and backward rate constants with the element in state C; *W*_R_(*n*) and *W*_T_(*n*) are the rate constants for changing the state from ‘R’ to ‘T’ and vice versa, respectively, in the *n*-th binding state. We further define

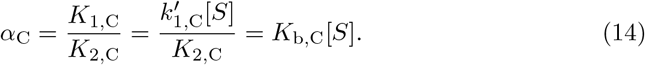

Here *α*_T_ = *cα*_R_ and 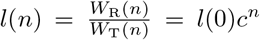. For significant cooperative behavior, *l* ≫ 1, *c* ≪ 1. The corresponding CMEs for the two states of this network element can 6 be written as

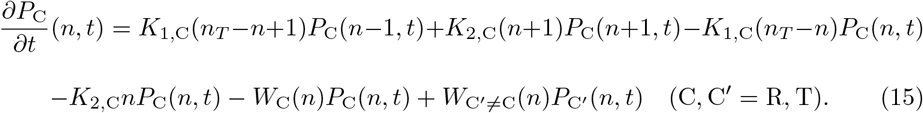

The equilibrium solution becomes

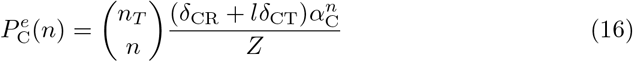

where 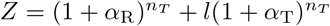. The mean binding number is given by

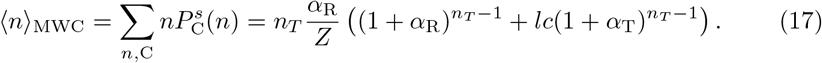

**Fig. 2.**
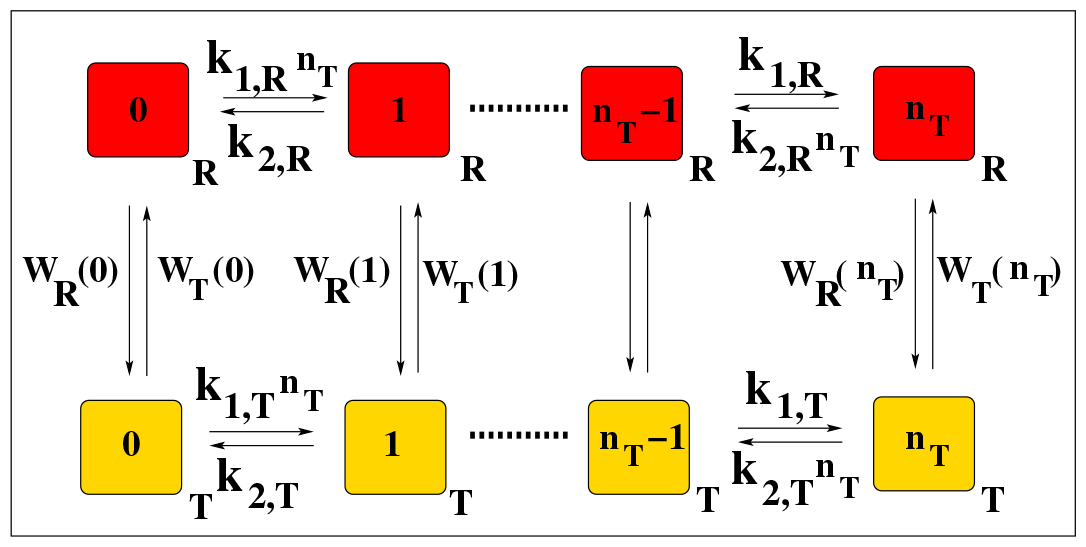
Schematic diagram of the MWC network. The squares denote states of the network element bound with a certain number of ligand molecules as indicated inside each square. The R and T subscripts stand for the state of the element of this network. The transition probabilities in forward and backward directions are also given (see text).

## 3 Shannon information and its relation with average binding and Hill slope for different cooperative networks

A. V. Hill proposed that if *n*_*T*_ number of ligand molecules bind to a multimeric protein simultaneously with dissociation rate, *K*_*d*_, the ratio of *θ* and (1 − *θ*) follows a monomial equation as[53]

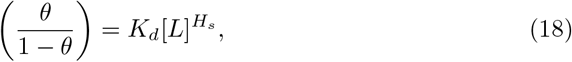

where fractional saturation, *θ* is defined as *θ* = ⟨*n*⟩*/n*_*T*_ and *H*_*s*_ is the Hill slope. Theoretically, the value *H*_*s*_ should be *H*_*s*_ = *n*_*T*_ but experimentally, it is observed in most of the cases that *H*_*s*_ < *n*_*T*_ as the ligand molecules do not bind to the protein simultaneously. Now, taking the logarithm of both sides of Eq.(18), we get the famous Hill equation[41, 53],

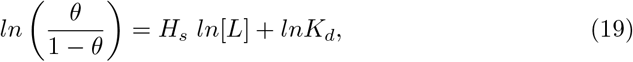

from which the Hill slope can be determined just by taking the derivative of Eq.(19) with respect to *ln*[*L*]. Thus, Hill-slope can be written in the form of logarithmic sensitivity [31, 40–43, 53]

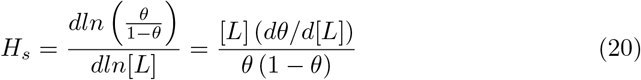

with the Hill-coefficient, *H*_*c*_ being defined as *H*_*c*_ = *H*_*s*_ |_*θ*=0.5_. Statistically, the Hill-slope is shown to be related to the variance of the binding number as[31, 43]

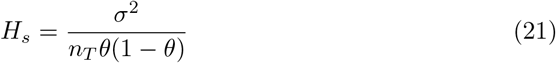

where the denominator in the right-hand-side is the variance of the corresponding non-cooperative case. The variance of the cooperative case becomes greater or less than that of the non-cooperative one for positive and negative cooperativity, respectively, with the mean binding number being same in both scenarios for a given ligand concentration[43].

Hill slope is the signature of the cooperative response of a system. This response is exhibited due to the transmission of information through successive binding of the ligand molecules. Here, our goal is to characterize the cooperativity in terms of the Shannon information associated with the states of the network with different binding numbers, *n*, varying from 0 (fully vacant) to *n*_*T*_ (fully occupied) in equilibrium. The Shannon information associated with state-n, denoted here by *I*_*n*_, is defined as[**?**]

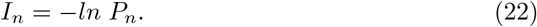

The expression of equilibrium probability distribution for all the cooperative networks studied here including the non-cooperative case can be written in the following generic form as a generalized Adair equation for stochastic binding

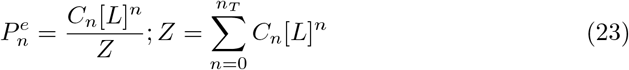

where *Z* depends parametrically on ligand concentartion [*L*].

We now study the variation of Shannon information with ligand concentration [*L*] in terms of the slope, denoted by *M*_*n*_, as

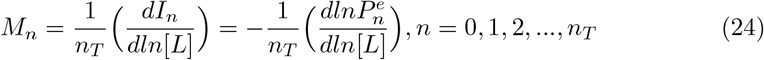

It follows from Eq.(23) and Eq.(24), that *M*_*n*_ has a generic form for all the cooperative networks given as

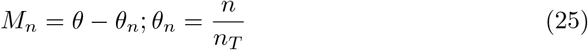

The details of the calculations for each individual network are given in the Appendix A. It follows from Eq.(25) that, the variation of Shannon information associated with the fully-vacant state (*n* = 0, *θ*_*n*_ = 0) and the fully-occupied state (*n* = *n*_*T*_, *θ*_*n*_ = 1) as a function of [*L*] are related to the fractional saturation of the cooperative binding network as

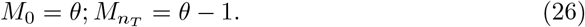

This clearly shows the connection of information with the binding statistics in terms of the average binding number; the latter is an experimentally measurable quantity.

Next, we explore further the logarithmic sensitivities of *M*_0_ and 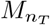 with respect to [*L*] and show that it also has a generic form for all the cooperative binding networks.

### 3.1 Logarithmic sensitivity of the slope of Shannon information

The logarithmic sensitivity of *M*_0_, denoted here by *ρ*_*n*_, is defined From Eq.(26) as

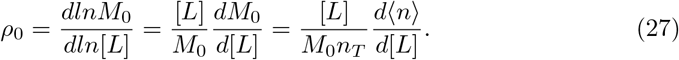

It is shown in Appendix B that, for all the networks considered here, one gets

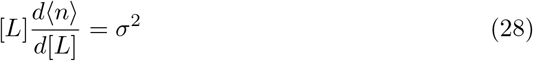

where *σ*^2^ is the variance of the binding number. Using Eq.(28) in Eq.(27), we get

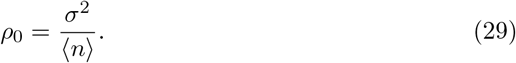

Now, it follows from Eq.(26) that 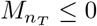 so, to derive the corresponding logarithmic sensitivity in this case, we take the negative of it, say 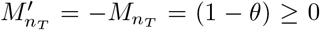. Then, for all the cooperative binding networks, one obtains

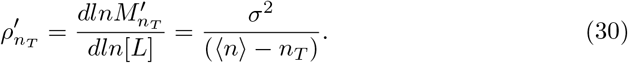

Interestingly, the logarithmic sensitivities in Eqs.(29) and (30) are of the form of Fano factor.

Finally, from Eq.(29) and Eq.(30) with Eq.(26), one can write

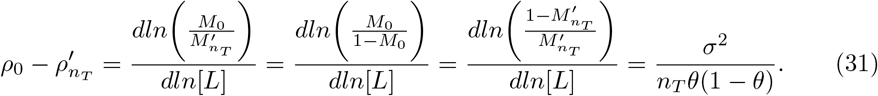

The right-hand-side of the above equation is, satisfactorily, the classical Hill slope given in Eq(21), where the denominator is the variance of a binomial distribution corresponding to the non-cooperative case (see Eq.(12)).

## 4 Numerical results

In this section, we perform some numerical experiments to verify the analytical findings. We take *K*_*b*_ = *K*_*b,R*_ = 0.1 *μM* ^−1^. For the KNF and the SP schemes, we consider an f-fold increase in each successive binding constant, *i*.*e*., *X*^(*j*)^ = *f*^*j*^*X*. Then the product is written as 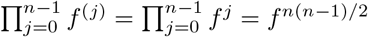. We take *f* = 1.5 for the numerics. To remind the reader, *f* = 1.0 gives the NC case. In case of the MWC model, we set *l* = 1000 and *c* = 0.001.

At first, we plot the fractional saturation *θ* as a function of [*L*] in Fig.3 for the different cooperative schemes (blue curves) along with *M*_0_ (red symbols) for *n*_*T*_ = 4.

**Fig. 3.**
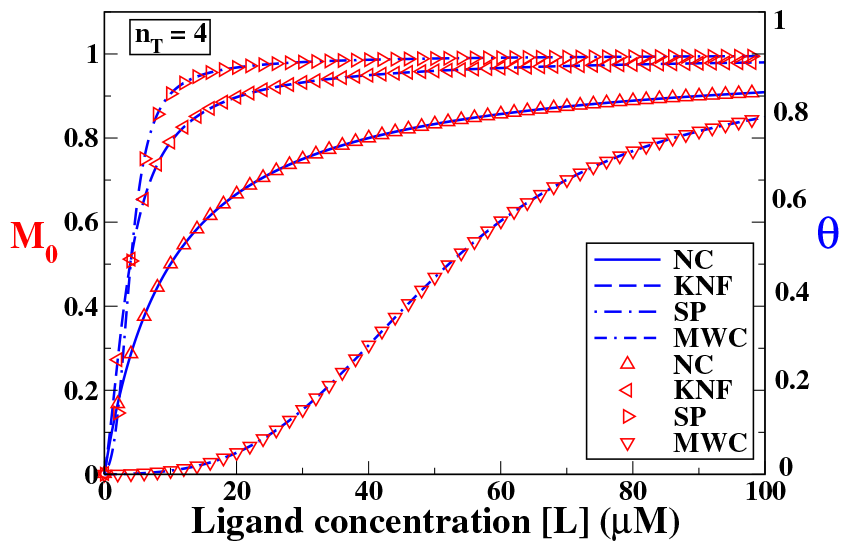
Variation of the fractional saturation *θ* as a function of [*L*] for the various networks of cooperativity with *n*_*T*_ = 4 (blue curves). The slope of Shannon information associated with the vacant state, *M*_0_, is also plotted (red triangles).

The latter are determined using numerical differentiation taking central difference. It is evident from the figure that the two quantities are the same as depicted in Eq.(26). For the given parameter values, the KNF and the SP scheme show steeper rise in the binding curve compared to the MWC model; the sigmoidal nature of the binding curve is more prominent in the latter.

Next, we show the variation of the logarithmic sensitivities, *ρ*_0_ and 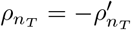 in Fig.4 as a function of [*L*] for all the schemes with *n*_*T*_ = 4. The Hill slope, generated from their combination as derived in Eq.(31), is also plotted (blue dashed curves). For the non-cooperative case (NC), *ρ*_0_ (black curve) and 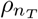 (red curve) vary monotonically in such a manner that the resulting Hill slope remains unity throughout (see Fig.4A). For all the other cooperative networks, the variations of *ρ*_0_ and 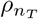 are non-monotonic as they pass through a maximum and generates a Hill slope that becomes greater than one(1) in intermediate ranges of ligand concentration, indicating positive cooperativity. The maximum of the Hill slope indicates the value of the Hill coefficient. Interestingly, *both ρ*_0_ *and* 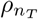 *become greater than unity for all the cooperative cases unlike the non-cooperative one where they remain less than or equal to unity*.

**Fig. 4.**
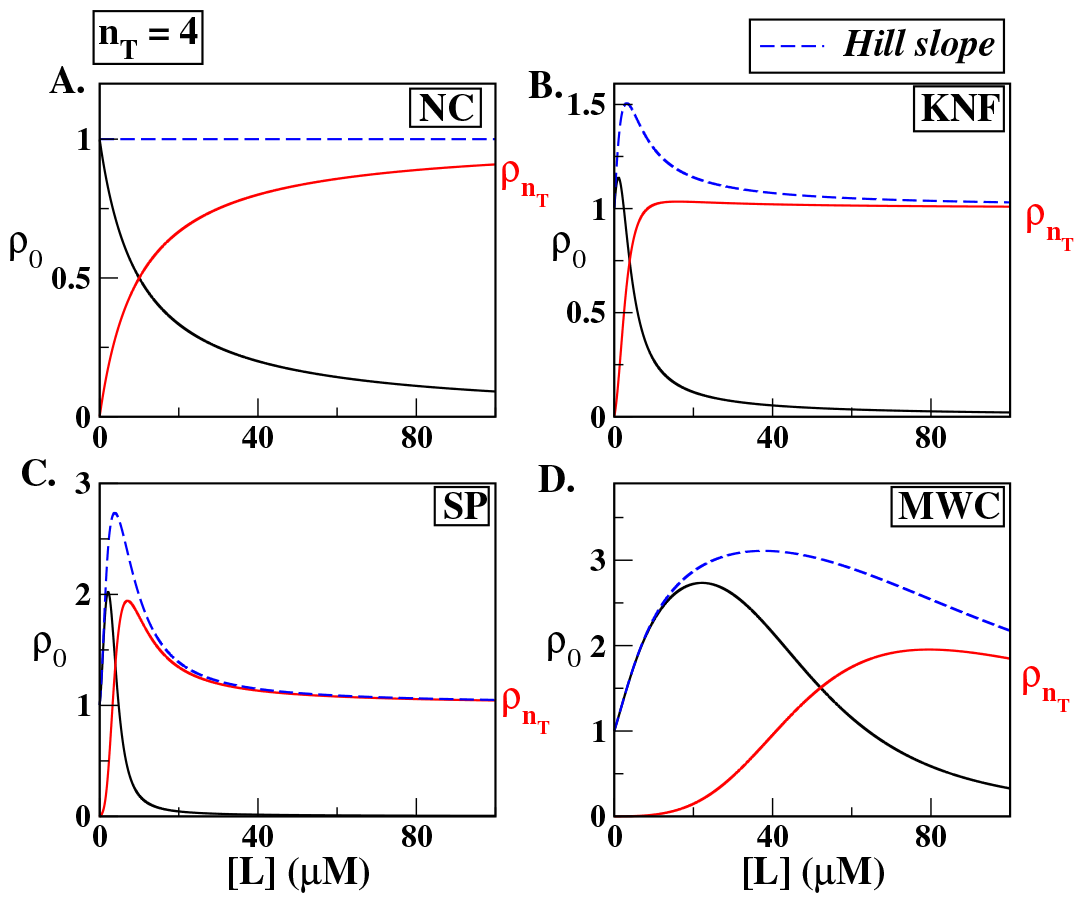
Variations of the logarithmic sensitivities, *ρ*_0_ (black curve) and 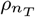 (red curve), as a function of [*L*] with *n*_*T*_ = 4 for the A. NC, B. KNF, C. SP and D. MWC networks. The Hill slope is also plotted (blue dashed curves).

## 5 Discussion and Conclusion

At cellular level, biological processes often exhibit cooperative response due to variation in the successive binding affinities of the ligands to the elements of their reaction networks. Generally, ligands, acting as signaling molecules, carry the cellular information which is transmitted to the elements of the network. Thus, a vital question arises: how we can characterize as well as measure the variation of the information transmission in the cooperative response of a reaction network? Here, we have addressed this question in terms of the variation of Shannon information associated with the state-probabilities of several well-known cooperative binding networks, *viz*., the KNF (as well as its variant) and the MWC network.

Cooperative response is generally characterized by measuring the value of the Hill-slope at the half saturation point *θ* = 0.5, known as the Hill-coefficient. Hill slope is derived from the Hill-equation that shows the dependence of a function of fractional saturation, *θ/*(1 − *θ*), on ligand concentration [*L*] as a logarithmic sensitivity. On a similar note, we have derived the slope of the Shannon information *M*_*n*_ associated with state-n as a function of *ln*[*L*]. It is shown that the slope has a generic form related to the fractional saturation *θ* for all the cooperative networks studied here. The slopes for the fully-vacant state (*n* = 0) and the fully-occupied state (*n* = *n*_*T*_), *M*_0_ and 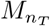, respectively, are given only in terms of *θ*. Specifically, *M*_0_ = *θ* and 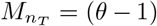. As *θ* is a measurable quantity in protein-ligand binding assays, the utility of this kind of relation regarding characterization and measurement of cooperative binding in terms of information is evident.

Further, the study of the logarithmic sensitivity, *ρ*_*n*_ of the slope *M*_*n*_ itself with respect to *ln*[*L*] reveals interesting information in terms of the ratio of variance to mean of the binding number with Fano factor like expressions. The combination of *ρ*_0_ and 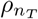 yields the traditional expression of Hill slope as the ratio of variances of the cooperative and non-cooperative cases.

In summary, our study characterizes variation of the Shannon information during the cooperative response by the networks due to binding of the ligands to the network elements at equilibrium. Theoretically, we characterize this variation by determining its slope at lower and upper ligand binding limits *i*.*e*., *n* = 0 and *n*_*T*_ which exhibits a generic form in terms of the mean binding number, an experimentally measurable quantity. At these limits, the slopes are independent of the nature of the network which illustrates a global behaviors about the information transmission processes due to ligand-binding. We also propose a methodology to determine these slopes at these extreme limits just by measuring the Hill slope. Our study provides an important insight in revealing the mechanistic details of the transmission process of the cellular information which is not generally obtained from the standard Hill-slope measurement. The study paves the path for further exploration of the information transmission processes at far from equilibrium that will be done in future.

## Appendix A Slope of the Shannon information for the various cooperative networks

### A.1 The KNF network

From Eq.(8) and Eq.(10), the probabilities of the fully-vacant and fully-occupied states can be written as

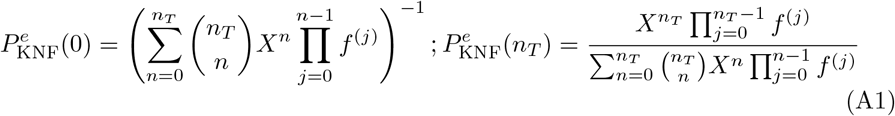

Then, the slope defined in Eq.(24) becomes

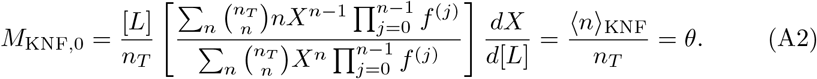

and following similar algebra, one gets

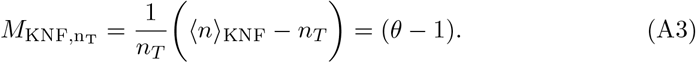

### A.2 The SP network

Using the equilibrium probability distribution, Eq.(9), for *n* = 0, we get from Eq.(24)

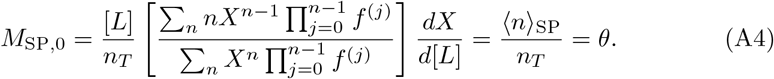

For *n* = *n*_*T*_, one can show in similar fashion that

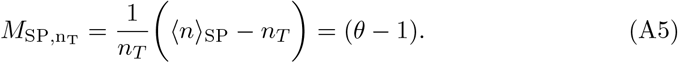

### A.3 The MWC network

In the case of this cooperative binding network, from Eq.(16) and Eq.(24), we obtain the expression of the corresponding slopes as

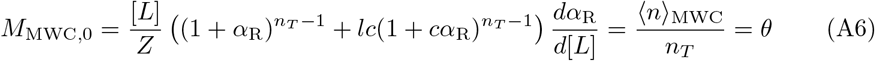

and

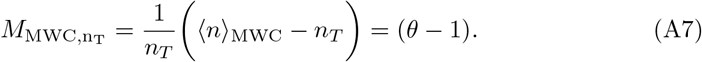

where we have also used Eq.(17).

Comparing the results obtained so far in this section, we can make the following observations: *For all the cooperative binding networks, the slope M*_*n*_, *n* = 0, *n*_*T*_ *can be expressed generally in terms of the fractional saturation as*

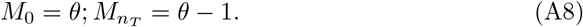

## Appendix B Slope of the average binding curve against ligand concentration

The slope 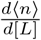 is an important feature of cooperative binding which is also intimately related to the Hill coefficient. Here, we derive the expression of this slope for the various cooperative networks.

For the KNF scheme, using Eq.(8), Eq.(10) and Eq.(11) along with the basic definition of mean binding, one can write

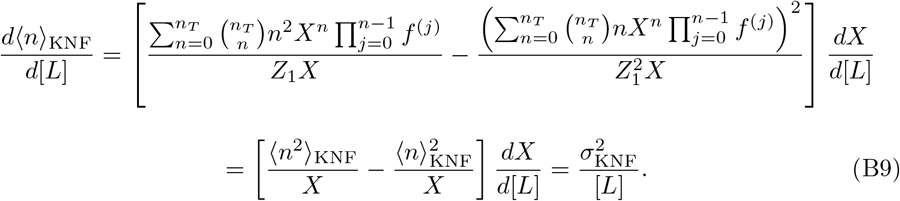

Here 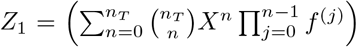 and 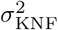 is the corresponding variance. It is obvious that a similar relation holds for the NC network which is a special case of the KNF network. Following similar procedure, one can easily check that Eq.(B9) is also valid for the SP cooperativity. Similarly, in the MWC network, from Eq.(16) and Eq.(17), we get

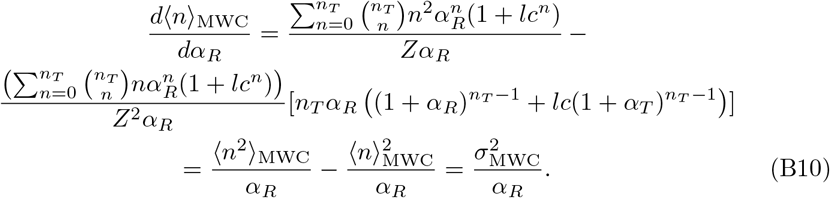

Using Eq.(14) with *C* = *R*, Eq.(B10) becomes

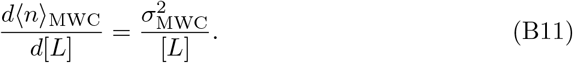

## Footnotes

1 Dedicated to Professor Gautam Gangopadhyay on his 60^th^ Birthday

